## Extended Data for "Implantable Bioelectronics for Gut Electrophysiology"

**The PDF file includes:**

Extended Data Figs. 1 to 10

**Other Extended Data for this manuscript include the following:**

Supplementary Movies S1 to S5

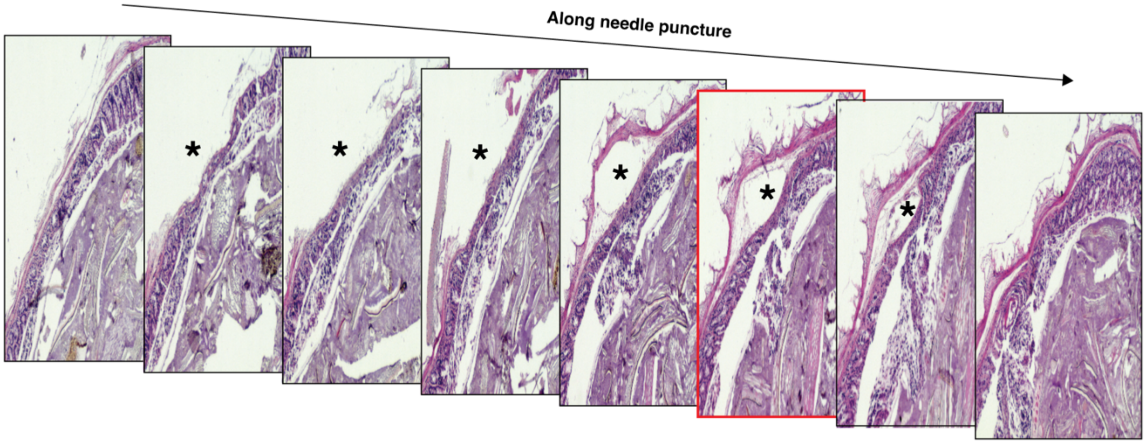


**Fig. 1. Histology showing the positioning of the tunnel used for implant placement.** Serial histological sections, stained with hematoxylin and eosin, showing the tunnel positioning for implant placement. The image outlined in red is the same as the image outlined in red in Fig. 1 of the main text.

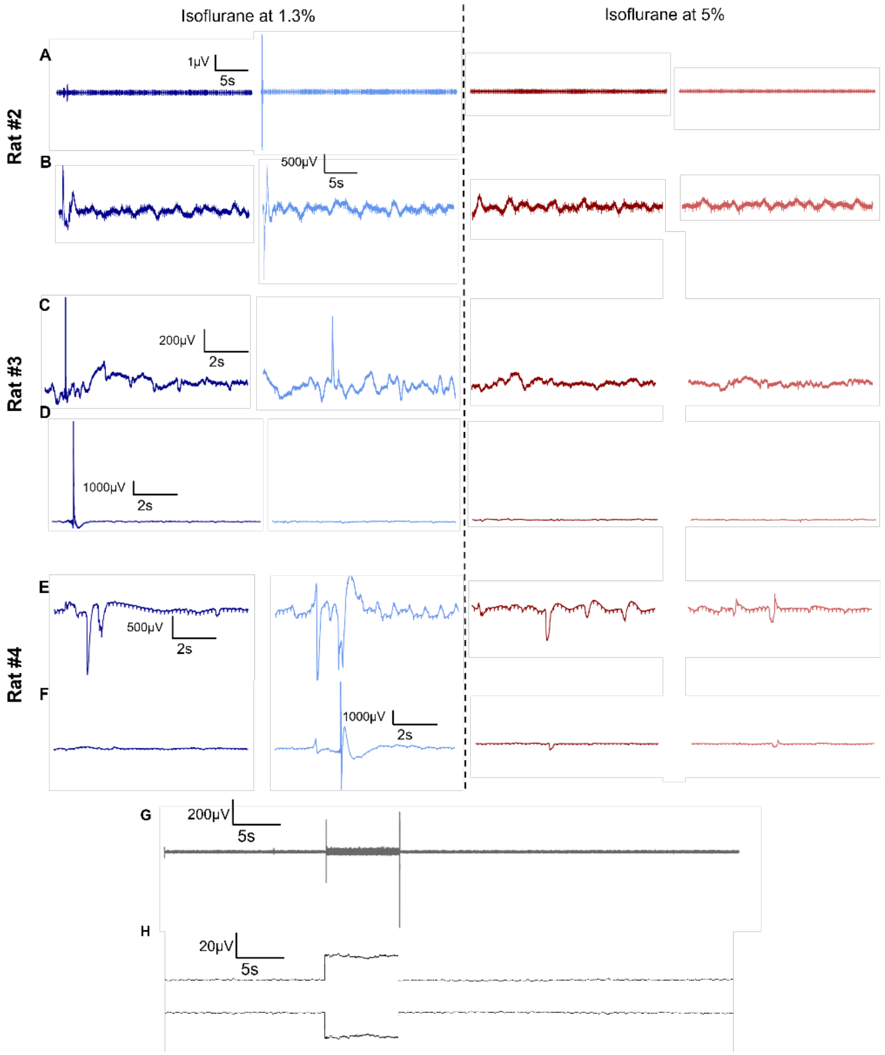


**Fig. 2. Additional instances of responses to mechanical stimuli under isoflurane and intraperitoneal urethane injection.** Under isoflurane: **A, C, E,** Extracted high-frequency bandpass traces corresponding to the distension segments**. B, D, F,** Extracted low pass traces corresponding to the distension segments. The extent and duration of the distension primarily relied on the level of ligation, carefully performed to close the lumen without causing tissue damage. The amplitude of the low and high pass voltage traces in response to the distensions was not uniform across all instances. For example, rats 2 and 4 show a larger response on the second distension compared to the first one. This variation is attributed to the inherent variability in pressure applied due to the placement of ligating sutures, etc. Given that the pressure levels were not identical, comparing response amplitudes directly should be avoided. Results for Rat#1 are shown in Fig. 2 in main manuscript. Under urethane: **G,** High-frequency filtered signal. High amplitude spikes correspond to mechanical artifacts due to insertion and removal of the needle **H,** Positive and negative envelopes of the signal. The portion of the colon with where the device was implanted was ligated and saline was injected manually using a syringe.

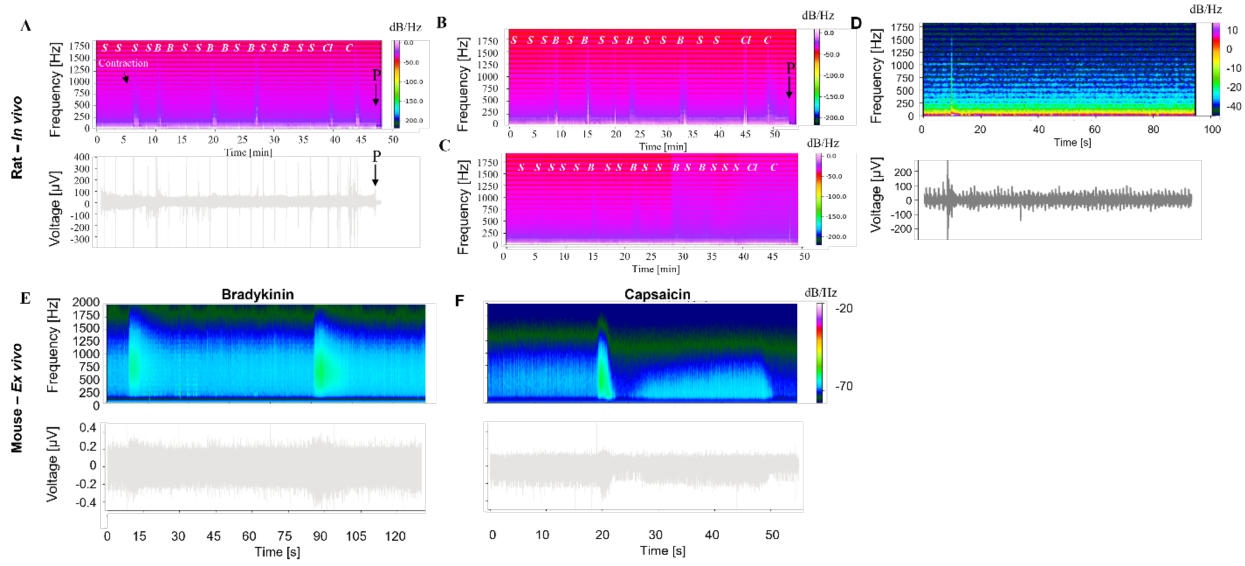

**Fig. 3. Full experimental traces of the ENS activation through drug administration. Rat *in vivo* models: A,** (top) Raw voltage signal corresponding to animal displayed in Fig. 2 in main manuscript showing high-amplitude peaks corresponding to drug administration and saline washes as conducted during the experiment. (bottom) Spectrogram (0-2000 Hz) from the same signal. Lettering shows times during which a substance was administered (S – Saline, B – bradykinin, $C_l$ – topical capsaicin, C – intraluminal capsaicin). The arrow on the far right of the trace and accompanying spectrogram depicts a final lethal injection of pentobarbital (P) into the heart, ending the experiment. This injection is followed by an immediate decrease in baseline, which is visible in the spectrogram. This particular trace also shows the contraction we observed, which is noted in Extended Data Fig. 5, and is denoted in the spectrogram. The spectrogram in Fig. 3a is the response at t = 10 minutes, Fig. 3b is the response at t = 39 minutes, and Fig. 3c is the response at t = 44 minutes. This same experiment was conducted in n = 6 rats, which is summarized in its entirety in Fig. 3d-f. Excluding the contraction, this spectrogram is representative for n = 4 of these rats (inclusive). For the remaining n = 2 rats, the timing and dosing differed slightly. These spectrograms are shown in **B** and **C**. **E,** Control recording: topical administration of saline. Raw voltage signal (top) and power spectra (0-2000Hz, bottom) of saline administration corresponding to same animal represented in Fig. 3 in main manuscript. ***Ex vivo* experiments on mouse distal colon:** Raw voltage signal and power spectra (0-2000Hz) of recordings from the lumbar splanchnic nerve (LSN) in response to bath infusions of **E,** bradykinin (1 μM) and **F,** capsaicin (500nM)

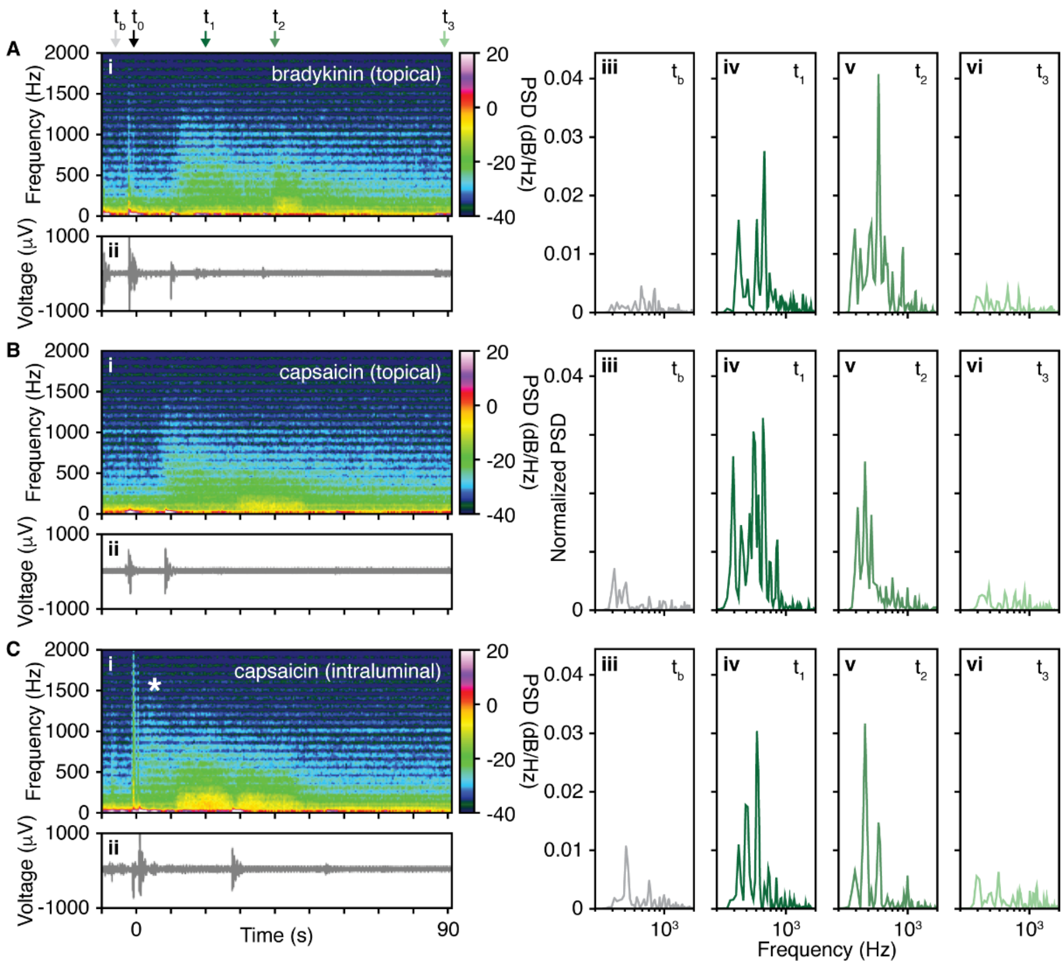


**Fig. 4 Temporal evolution of frequency responses to drug administration (i)** normalized power spectra (zoom in 200 - 1400 Hz) and **(ii)** raw voltage signal as represented in Fig. 3 in main text. (**iii**) $t_b$, (**iv**) $t_1$, (**v**) $t_2$, and (**vi**) $t_3$, time points semi-arbitrarily chosen to highlight change in the relative power spectra across the recording and indicated in **A, B** and **C** with color-coded, labeled arrows. Drug administration occurred at approximately $t_0$ for **A**, **B**, and **C**.

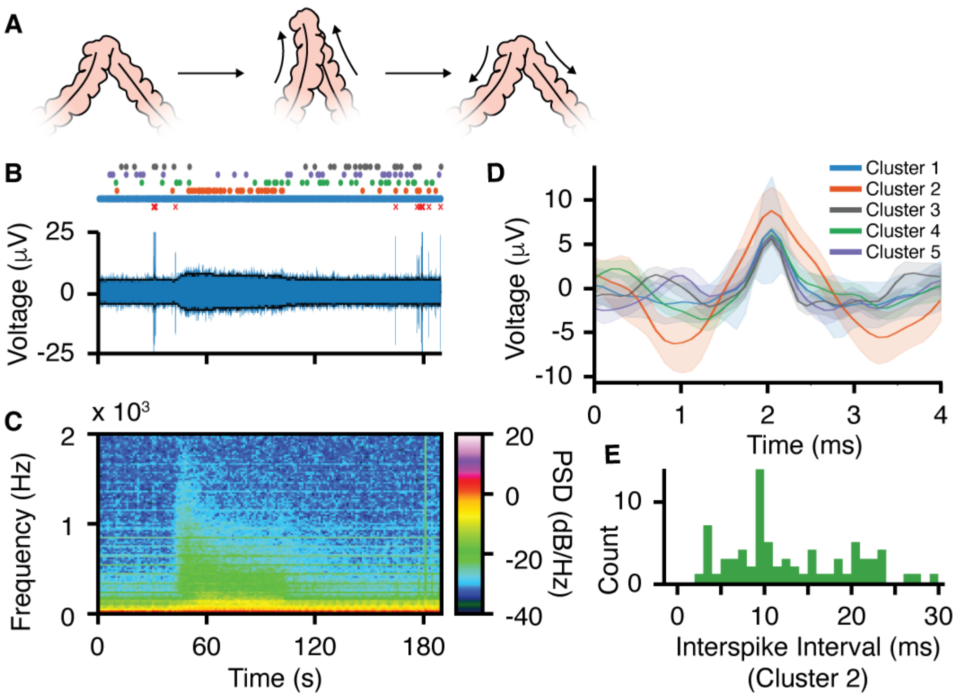


**Fig. 5 Recording of Colonic Contraction. A**, Schematic illustrating the approximate contractile conformation that was observed. **B**, Raw temporal signal (n=1) with localisation of detected spikes (circles) and artifacts (red crosses) corresponding to each cluster in **D.** Sharp signal is the saline addition. **C,** Temporally aligned spectrogram of the same contraction trace. **D,** Average waveforms of identified clusters from detected spikes. **E,** Inter-spike-interval (ISI) plot showing distribution of spikes corresponding to cluster 2. The wide distribution of inter-spike intervals (ISIs), with a primary peak at ~9 ms and no clear harmonic structure, supports the neural origin of the signal.

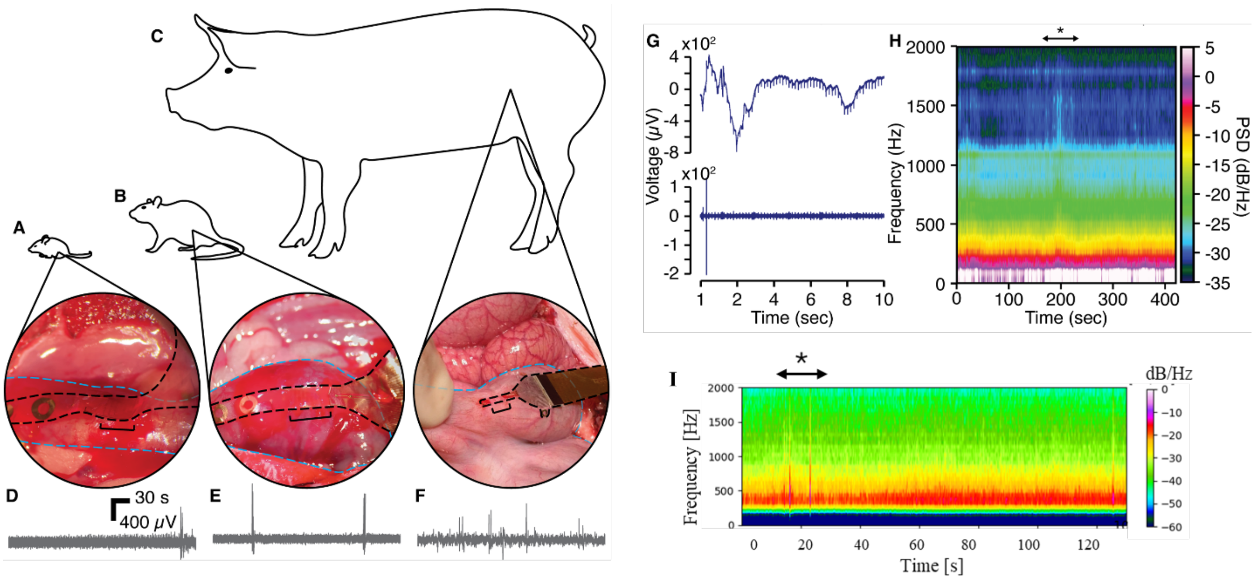


**Fig. 6. Surgical placement and electrophysiological recording demonstration in mouse and pig.** Image showing surgical placement of implantable device in **A** mouse colon, **B** rat colon, and **C** pig colon. For each image, the device is outlined with black dotted lines, and the colon is outlined with blue dotted lines. The black bracket indicates the portion of the device that resides within the colonic wall. Representative voltage traces, bandpass filtered between 0 and 2000 Hz, for **D** mouse, **E** rat, and **F** pig. We see similar signal-to-noise ratios for each system. **G** Distension trace for mouse showing time-synced (top) low-pass (0-300 Hz) and (bottom) high-band pass (300-2000 Hz). **H** Spectrogram showing response to capsaicin administration in the pig colon. For this experiment, ~10 mL of capsaicin was poured topically onto the region containing the implant over the course of 1-minute. This spectrogram begins (time at 0 seconds) at the culmination of the capsaicin dosing. The maximal visible response is highlighted using an asterisk. **I** Topical administration of saline – control experiment. Power spectra (0-2000Hz) of saline administration corresponding to same animal represented in H. These pilot experiments were performed in a similar fashion to the rat data, although further refinement may be necessary if conducting an entire study in a different species. However, given the conservation in neuronal density across vertebrate species [46,47], the data we collect should be of similar origin to those data from the rat.

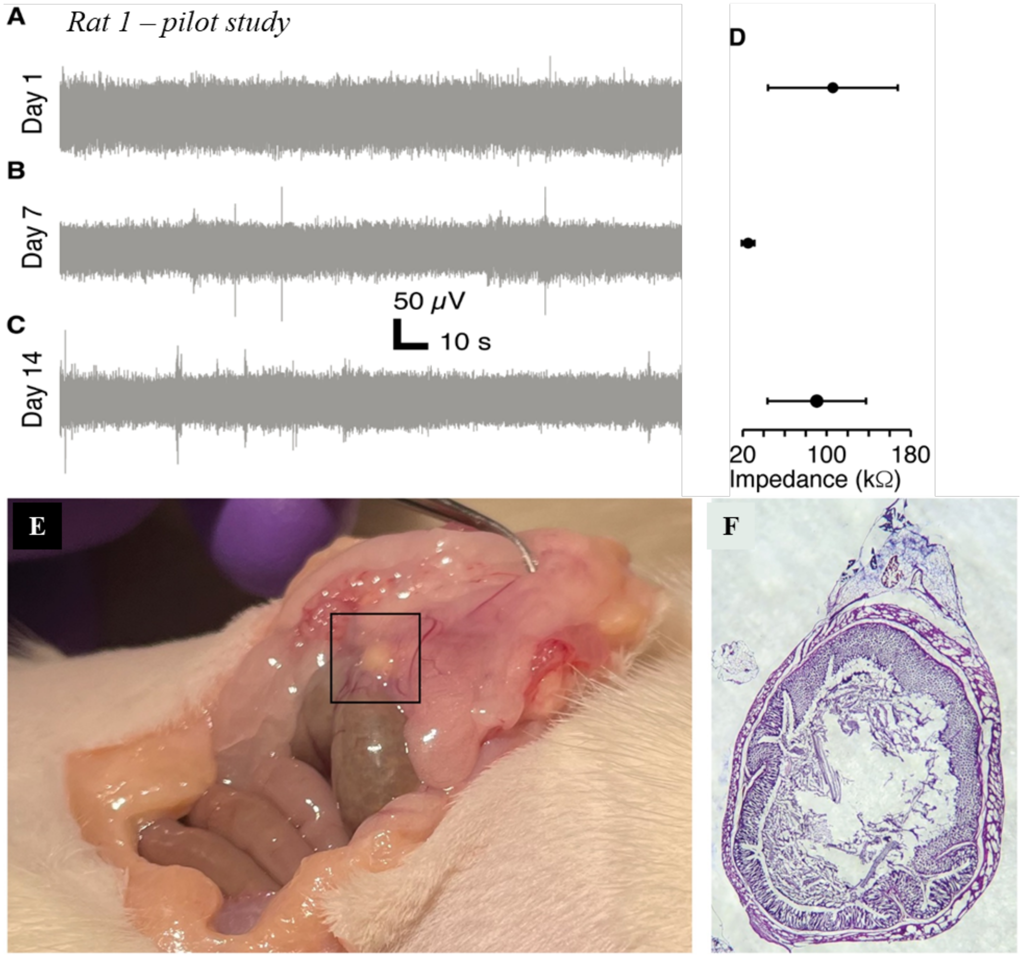


**Fig. 7. Feasibility results from pilot study. A-C,** Representative traces of representative rat 1 in pilot study showing chronic implant stability over 2 weeks at **A,** Day 1, **B,** Day 7, and **C**, Day 14. Traces were filtered using a high band pass filter from 300 to 4000 Hz. **D,** Impedance values of all available channels for each day. **E**, Post-mortem and histology from chronic implant placement. Fibrosis was noted onto the colon from the peritoneal wall, beginning at the point where the implant crosses into the peritoneal cavity. This region is highlighted by a black box in the image. **F,** Hematoxylin & eosin stain of colonic section in the vicinity of the implant, showing intact colonic wall. Exact identification of implant positioning along the axis of the colon was not achieved during sectioning.

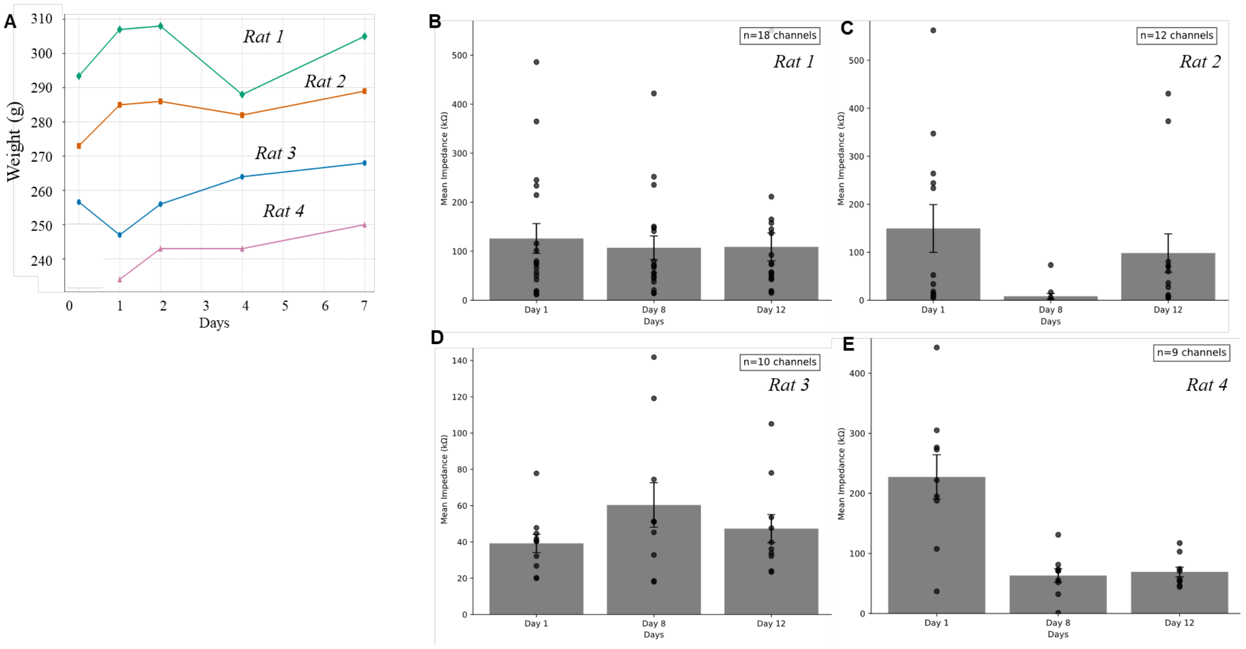


**Fig. 8. Stability and health metrics during behavioural recordings (n = 4). A,** Rat weight evolution during the first week. For the first 7 days, we monitored the weight evolution of the rats. A slight decrease was observed over the first four days, which is expected following surgical intervention. However, all animals recovered and surpassed their initial weight within a week, indicating good tolerance to the implant. **B-E,** Mean impedance values for each day with standard deviation for the same active channels across Days 1, 8, and 12 for rats 3 to 6 (**B to E**, respectively).

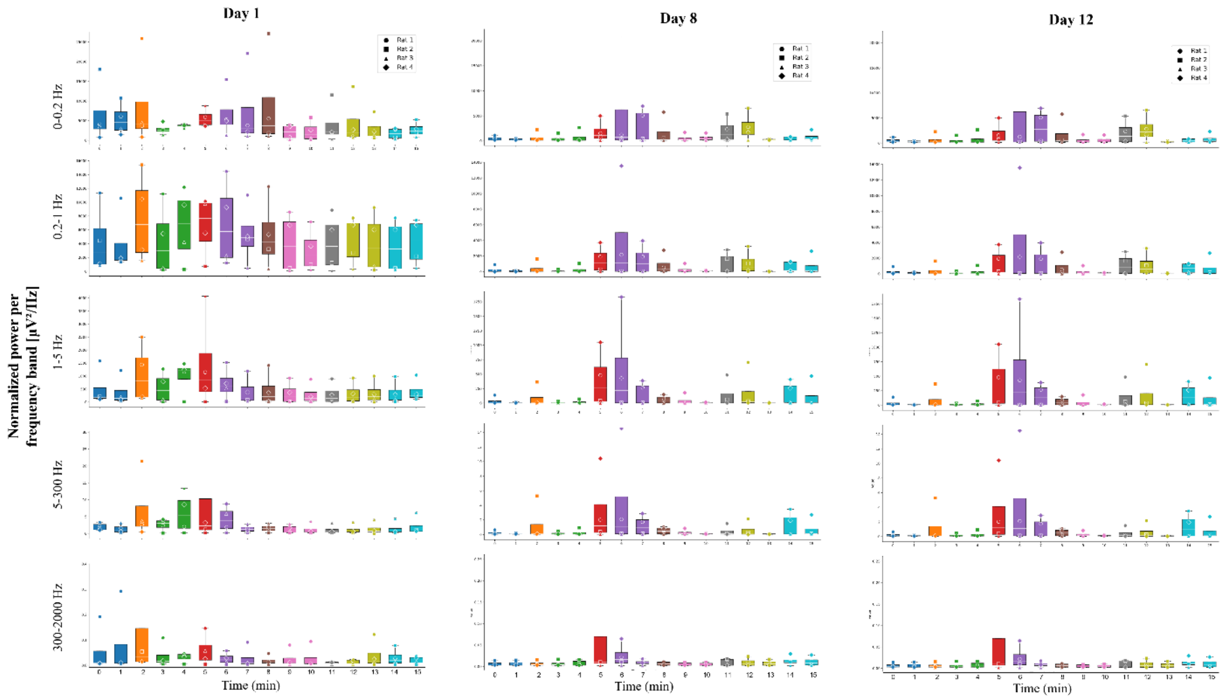


**Fig.9. Boxplots showing the normalized power per frequency range response of all rats (n = 4) in the 15 minutes of open field test on days 1, 8 and 12.** For each rat, raw signals were referenced to the mean signal across all valid channels. These referenced signals were then segmented into 1-minute intervals to analyze the temporal evolution of the signals. Within each interval, we divided the data into predefined frequency bands using bandpass filters for detailed examination. The power within each frequency band was normalized by the bandwidth to allow for fair comparison across bands of different widths (normalized power per frequency range). **Day 1**: A first increase in the response is evident within 3 min in all bands, with a secondary peak at 5 to 8 minutes depending on the frequency band. **Day 8:** a smaller first increase in the response is present within 3 min in all bands (note the different y-scale), with a secondary peak at 5 to 8 minutes depending on the frequency band. **Day 12:** a smaller first increase in the response is present within 3 min in all bands (note the different y-scale), with a secondary peak at 5 to 8 minutes depending on the frequency band. The y-scale has been optimized per day to help visualizing the smallest differences in day 12 compared to day 1.

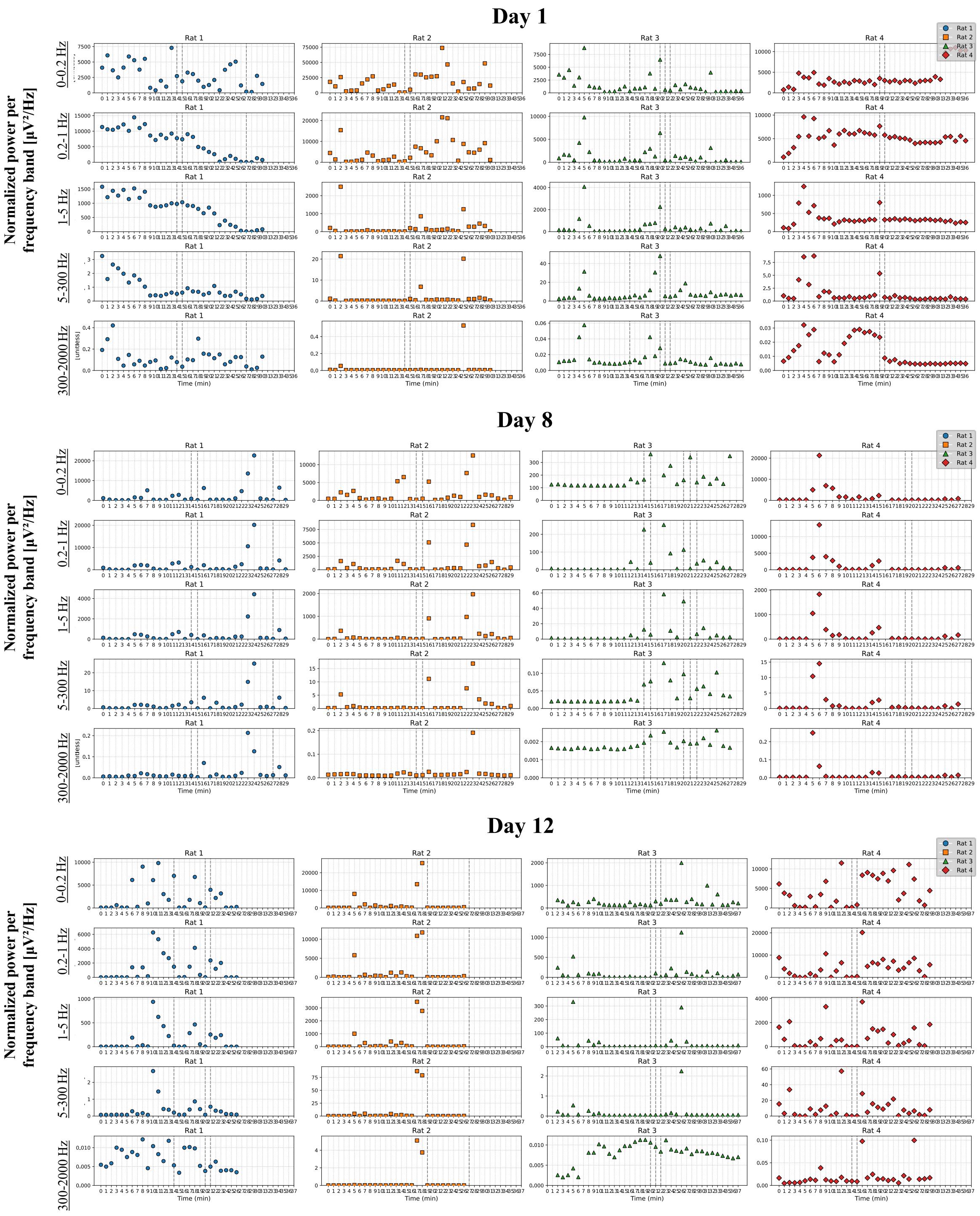


**Fig. 10. Scatter plots showing the temporal evolution of the normalized power per frequency band for the duration of the chronic recording for each rat on Days 1, 8, 12.** Dotted vertical lines depict the times when the rats were eating. Note that the amount of food differs at each feeding event, as rats were eating as desired. This causes a wide range of response amplitudes at each feeding instance.

**Supplementary Movies 1 and 2. Videos of *in vivo* placement of implant under anesthesia.** Videos are looped to allow for examination for fine movements of tissue.

**Supplementary Movie 3. Mechanical distension of ligated colon section through intraluminal saline injections.** The implant conforms to the expansion of gut tissue.

**Supplementary Movie 4. Freely-moving animal during chronic recording.**

**Supplementary Movie 5. Freely-moving rat, shortly after placement into open-field environment.**
